# *Staphylococcus aureus* derived extracellular vesicles trigger multiple inflammatory pathways in host cells and deliver their RNA cargo following their internalization and lysis within late endosomes

**DOI:** 10.1101/2025.10.06.680696

**Authors:** Julia Papail, Nathalie Daniel, Ligia Prado, Daniele Vassaux, Sandrine Péron, Céline Even, Véronique Lebret, Brenda Silva Rosa Da Luz, Yann Le Gouar, Julien Jardin, Siomar de Castro Soares, Ken-Ishi Yoshida, Vasco Azevedo, Svetlana Chabelskaya, Yves Le Loir, Éric Guédon

## Abstract

*Staphylococcus aureus* is a major opportunistic Gram-positive pathogen whose virulence can be partly mediated by extracellular vesicles (EVs). These nanosized particles carry a diverse cargo and can play key roles in host-pathogen interactions. Here, we investigated the production, composition, and host cell interactions of EVs secreted by the methicillin-resistant *S. aureus* strain N315, one of the most widely studied clinical isolates. To mimic infection, EVs produced from RPMI supplemented with 10% LB were purified by size exclusion chromatography, and characterized by nanoparticle tracking analysis and electron microscopy. N315 EVs contained a diverse molecular cargo, selectively enriched, and including proteins, lipoproteins, DNA, RNA, lipoteichoic acid, and peptidoglycan. Protection assays and confocal microscopy confirmed the intravesicular localization of RNAs and proteins. N315 EVs were recognized by multiple host pattern recognition receptors (PPRs) in human MG-63 cells, located at the cell surface (TLR1, TLR2, TLR4, TLR6), within endosomes (TLR3, TLR7, TLR4) and in the cytoplasm (NOD2). EV exposure promoted the selective upregulation of these PRRs, activated the NF-κB and JAK/STAT signaling pathways, and induced the expression of pro-inflammatory cytokines such as IL1β, IL6, and CXCL8. N315 EVs were internalized via a dynamin-dependent endocytosis mechanism and trafficked to late endosomes, where RNA cargo release occurred. Importantly, inhibition of EVs uptake altered cytokine gene expression, confirming that internalization is required for immunomodulation. This work provides a better understanding of the mechanisms of interaction of *S. aureus* EVs with host cells and sheds light on the fate of EVs and their cargo once internalized by cells.

**IMPORTANCE:** Antibiotic resistance, coupled with the limited discovery of new antimicrobial agents, poses a growing threat to public health. *Staphylococcus aureus* is one of the six highly virulent and antibiotic-resistant bacterial species responsible for both severe hospital- and community-acquired infections. This pathogen demonstrates a remarkable ability to adapt to diverse hosts and environments and to produce a broad arsenal of virulence factors. Gaining deeper insight into its pathogenic mechanisms and host interactions may reveal novel therapeutic targets and alternative treatment strategies. Over the past decade, a key advancement has been the identification of extracellular vesicle production by *S. aureus* and its critical role as a new secretion system for virulence factors. This study shed light on the immunogenic effect of EVs and their role as a vehicle for the delivery of their contents into host cells. These adjuvant-like properties open new avenues for research and potential intervention.

## INTRODUCTION

*Staphylococcus aureus* is one of the most common bacterial species found in human and veterinary medicine. It is an opportunistic Gram-positive pathogen primarily associated with warm-blooded animals. In humans, 30% of the population are asymptomatic carriers of this bacterium, with colonization mainly on the skin and in the nasal mucosa (1). However, under certain conditions of dysbiosis, such as a weakening of the host’s immune defenses or the presence of wounds, *S. aureus* can cause nosocomial and community-acquired diseases, ranging from mild with suppurative skin infections to severe and invasive infections (2–4). Managing *S. aureus* infections has become increasingly difficult due to its strong adaptive capacity and stress tolerance (5–7). The intensive use of antibiotics has further promoted the selection of resistant and multidrug-resistant strains, notably methicillin-resistant *S. aureus* (MRSA) strains, which are associated with significant morbidity, mortality, prolonged hospital stays, and high healthcare costs (8). These challenges emphasize the urgent need to develop alternative strategies to fight this pathogen, which requires a better understanding of its virulence mechanisms and interactions with the host.

The ability of *S. aureus* to colonize different hosts and cause a wide range of infections relies on its production of numerous virulence factors, such as surface-exposed and secreted proteins, both responsible for the colonization and the disease severity (9, 10). These virulence determinants are highly regulated and allow *S. aureus* to switch between a commensal and pathogenic lifestyle depending on host context (11). As described for several other pathogens, it has recently been proposed that extracellular vesicles (EVs) produced by *S. aureus* may constitute an integral component of its virulence arsenal (12, 13). Bacterial EVs are nanosized, lipid bilayer-enclosed particles. They typically range from 30 to 300 nm in diameter and enclose a complex molecular cargo composed of proteins, lipids, RNA, DNA, and metabolites (14–17, 13). By mediating the transfer of bioactive components, EVs contribute to multiple aspects of bacterial physiology, such as stress adaptation, intercellular competition, lateral gene transfer, biofilm development and antibiotic resistance (18–21). Numerous studies, notably on *S. aureus*, have also underlined their role in intra- and inter-species communication, and in host-pathogen relationships (12, 13). EVs secreted by diverse *S. aureus* strains, of both human and animal origin, contribute to pathogenesis in playing roles in the dissemination of virulence factors, and modulation of host immune responses (22–27).

Although EV production and cargo composition can vary between strains, conserved features emerge underlining common functional roles in *S. aureus* biology. In particular, the protein cargo of EVs is enriched in lipoproteins and virulence factors, and includes a conserved core proteome (17). For instance, the presence of pore-forming toxins inside EVs may account for their cytotoxic effects on phagocytic cell lines (neutrophils and macrophages), as well as on erythrocytes and epithelial cells (28–30, 27). Hong *et al*. (2014) demonstrated that the severity of atopic dermatitis caused by *S. aureus* was exacerbated in the presence of EV-associated α-hemolysin compared with its soluble form. Hemolysin associated with EVs was found to be much more cytotoxic for keratinocytes (31). In addition to disrupting the skin barrier, only the EV-bound hemolysin form induces local skin inflammation mediated by neutrophiles. These observations indicate EV-associated cargo can be biologically active and suggest that EVs not only serve as protective carriers but also as efficient delivery systems for virulence factors.

In addition, *S. aureus* EV cargo is composed of pathogen-associated molecular patterns (PAMPs) such as lipoproteins, peptidoglycan (PG), and nucleic acids, which can be recognized by host pattern recognition receptors (PRRs), including Toll-like and NOD-like receptors, located at the cell surface, within endosomes and in the cytosol. Their activation triggers downstream signaling cascades, leading to cytokine and chemokine production, immune cell recruitment, and inflammation. The ability of *S. aureus* EVs to activate the NF-κB pathway and induce the secretion of pro-inflammatory cytokines has been extensively documented in recent years (23, 28, 32, 29, 33, 27). For instance, Tartaglia *et al*. (2018) and Kopparapu *et al*. (2021) reported that EVs derived from both animal and human isolates of *S. aureus* stimulate the secretion of pro-inflammatory cytokines not only *in vitro* but also *in vivo* (34, 26). Many studies agree on the crucial role of TRL2, a surface PRR recognizing bacterial lipoproteins, in mediating NF-κB activation in response to *S. aureus* EVs (34, 35, 27). More recently, the involvement of additional PRRs has also been demonstrated (23). Despite these observations, the mechanisms by which *S. aureus* EVs interact with host cells, the fate of their cargo, and their contribution to host immune modulation remain incompletely understood.

This study aims to provide new insights into the interactions between *S. aureus* EVs and host cells, focusing in particular on the nature and fate of their cargo once internalized. First, we characterized the production, the molecular content, and the immunomodulatory effects of EVs secreted by N315, a MRSA human strain of *S. aureus*. We showed that N315 EVs contain a diverse molecular cargo, notably numerous PAMPs, and demonstrated selective packaging and protection of proteins and RNAs. Next, we showed that N315 EVs are detected by a variety of PRRs on human osteoblast-like MG-63 cells and activate key signaling pathways, including NF-κB, JAK/STAT and inflammasome. Finally, we demonstrated that EVs are internalized by host cells via dynamin-dependent endocytosis, release their RNA cargo in late endosomes, and modulate host gene expression in an internalization-dependent manner. By contributing to a better characterization of EVs, this study provides a better understanding of the interactions between *S. aureus* and its host.

## MATERIALS AND METHODS

### Bacterial strains, and growth conditions

The methicillin-resistant *S. aureus* strain N315 (CIP 107652), isolated in 1982 from a pharyngeal smear of a patient in Japan, was grown in eukaryotic cell culture medium RPMI (Sigma, R8758) supplemented with 10% of LB (Luria Bertani Broth: Tryptone 10 g/L, Yeast Extract 5 g/L, NaCl 5 g/L) (*i.e.*, RPMI+LB) at 37°C with shaking at 180 rpm (36). *S. aureus* RN4220 was cultivated in brain heart infusion broth (BHI; pH 7.4; Becton Dickinson, Le Pont de Claix, France). *Escherichia coli* NEB® 5alpha (New England Biolabs) and *Bacillus subtilis* KT01 (37, 38) were cultivated under vigorous shaking in LB medium at 37°C. When required, antibiotics (Sigma-Aldrich) were added to the media at the following concentrations: ampicillin (100 μg/mL for *E. coli),* spectinomycin (100 μg/mL for *B. subtilis*), and chloramphenicol (10 μg/mL for *S. aureus*). The colony-forming unit (CFU)/mL was determined after 24 h of incubation at 37 °C on BHI or LB agar.

### Construction of translational sfGFP fusion with the *S. aureus* Eno protein

To illustrate specific protein packaging into EVs, we selected enolase (Eno) as a marker of EV loading. Eno has been previously identified as part of the core EV proteome of many Gram-positive bacteria (17) and was detected here in N315 EVs. For that N315 was engineered to express a translational fusion between Eno and the superfolder green fluorescent protein (sfGFP) on a replicative vector. Primers used are listed in Table S1. The *eno* and *sfGFP* genes were amplified by PCR using Q5 Hot Start High-Fidelity DNA Polymerase (M0493, New England Biolabs) from genomic DNA of *S. aureus* N315 and *Bacillus subtilis* KT01, respectively. The latter strain carried a stable chromosomally integrated single copy of the *sfgfp* gene. Genomic DNA was recovered using DNeasy Blood and Tissue kit (69504, Qiagen). PCR fragments were purified with NucleoSpin® Gel and PCR Clean-up kit (740609, Macherey-Nagel), and assembled using the Gibson assembly method (Gibson Assembly Cloning kit, E5510, New England Biolabs) (39). The resulting *eno*-*sfGFP* translational fusion amplicon was restricted by *Sph*I and *Bam*HI and cloned into the *Sph*I/*Bam*HI digested pCN35c vector (40, 41). The resulting multicopy plasmid pEV1 was transformed into the RN4220 *S. aureus* strain, and then in *S. aureus* N315 by electroporation (2.45 kV, 25 μF, 100 Ω; BIO-RAD, Gene Pulser Xcell) to give strain GMS0045 (N135 pVE1). Plasmid preparations were performed using NucleoSpin Plasmid kit (740588, Macherey-Nagel) according to the manufacturer’s recommendations. We verified the sequence of the recombinant pEV1 plasmid from strain GMS0045 by long-read sequencing (ONT Lite Whole plasmid sequencing, Eurofins Genomics).

### Extracellular vesicles isolation

N315 EVs were isolated from bacterial culture supernatants inoculated with an 8 h preculture (0.5%) in 250 mL of RPMI+LB and incubated at 37°C with shaking at 180 rpm for 16 h, until the stationary phase (OD_600nm_ > 2.0, ∼ 10^9^ CFU/mL). Bacterial cell cultures were centrifuged at 3,100 × *g* for 30 min and filtered through 0.22 μm Nalgene top filters (Thermo Scientific). For routine experience, 1 L of culture supernatant was used, obtained by pooling four 250 mL cultures at this step. The 1 L filtered culture supernatant was concentrated with Amicon Ultra-15 centrifugal filter units (100 kDa molecular weight cut-off, Millipore) by successive centrifugations (3,220 x *g*). The concentrated suspension of EVs was recovered in PBS buffer (D8537, Sigma-Aldrich) and further purified by size exclusion chromatography (SEC) using qEVoriginal 70 nm columns (Izon) as recommended in the manufacturer’s instructions. Fractions enriched in EVs and almost free of contaminants were pooled (fractions 2 to 6) and concentrated with Amicon Ultra-15 centrifugal filter units with 100 kDa cut-off. The recovered EVs were analyzed by Qubit (Invitrogen, Qubit 4 fluorometer) for DNA and RNA content and with Pierce^TM^ BCA Protein Assay Kits (BCA) for protein concentration. For the choice of pooled fractions, each 400 µL fractions were collected sequentially and analyzed for particle content and protein contaminants (see Fig. S2 for more details on SEC fractions).

### Determination of EVs Sizes and Concentrations

Nanoparticle Tracking Analysis (NTA) using an sCMOS camera and a Blue488 laser (Nano Sight NS300) was performed to assess EVs size and concentration. Samples were diluted into PBS to achieve an optimal concentration of around 40 to 100 particles per frame and submitted to a constant flux generated by a syringe pump (speed 50), at 25°C. Results were retrieved from 5 × 60 s videos recorded with camera level at 15-16 and threshold at 3-5, while other parameters were adjusted as necessary.

### Transmission electron microscopy imaging

For microscopy imaging, EVs were isolated by SEC using TBS buffer (150 mM NaCl; 50 mM Tris-Cl, pH 7.5) instead of PBS. Copper EM grips (Electron Microscope Sciences FF200-Cu) were first exposed to a glow discharge of 20 mA. Ten μL of a solution containing around 10^11^ particles per mL were fixed with 4% paraformaldehyde (PFA) and placed on glow-discharged grids for 30 s. After removing of liquid excess with filter paper, grids were stained with 2% uranyl acetate and observed with JEOL 1400 transmission electron microscope operating at 80-120 kV.

### Nuclease protection assays and RNA purification

The procedure used for nuclease protection assays was also used to remove external RNA contaminants before EV RNA extraction. To degrade contaminant RNAs potentially protected by proteins from RNase A digestion, 5.0 × 10^10^ EVs were first treated with proteinase K (50 µg/mL, 03115836001, Sigma), followed by inhibition of protease activity with 5 mM phenylmethylsulfonyl fluoride (PMSF) (93482, Sigma). RNase A (0.5 µg/mL, 19101, Qiagen) was then added for 5 min at 25 °C, after which RiboLock RNase inhibitor (EO0381, Thermo Scientific) was applied according to the manufacturer’s instructions. For RNA extraction, EVs lysis was performed by mixing the samples (v/v) with 350 µL of lysis buffer (0.5% SDS w/v, 30 mM sodium acetate, 1 mM EDTA, acid-buffered at pH 5.0), glass beads and 700 µL of acid phenol, followed by mechanical disruption for 30 s in Precellys homogenizer at 6,500 rpm. Samples were then centrifuged (16,000 x *g*, 4°C, 5 min) and the upper phase was transferred to a new tube, mixed with an equal volume of phenol and centrifuged under the same conditions. The resulting upper phase was recovered, mixed (v/v) with chloroform:isoamyl alcohol 24:1, and centrifuged (16,000 x *g*, 4°C, 5 min). The final upper phase was collected, supplemented with 0.2 µg/µL of glycogen (R0551, Thermo Scientific), 10% volume of NaAc 3M and 2.5 volumes of ice-cold 100% ethanol, and stored overnight at −20°C. Samples were then centrifuged (16,000 x *g*, 30 min, 4°C), and the pellets were washed with 1 mL of cold 70% ethanol. Finally, the pellets were dried with a SpeedVac concentrator and dissolved in RNase-free water. RNAs were cleaned up and treated with DNase I using the RNA Clean and Concentrator^TM^ kit (Zymo Research). Samples were kept at −80°C until use. Samples were analyzed by Nano Drop (Labtech, Spectrophotometre Nanodrop ND1000), Qubit (Invitrogen, Qubit 4 fluorometer) and Bioanalyzer. This latter analysis was realized on RNA 6000 Pico Kit by the company Helixio, Saint-Beauzire, France.

### Protease protection assays

Proteinase protection assays were performed with 2.0 × 10^10^ EVs (equivalent to 10 µg proteins) per condition. Five conditions were tested: i) untreated EVs (control), ii) EVs incubated with PMSF (5 mM, 10 min, RT), iii) EVs treated with radioimmunoprecipitation assay buffer (RIPA; 1X final; 30 min, 4°C, 2,000 rpm in thermomixer) to permeabilize EVs and then with proteinase K (50 µg/mL, 10 min, 37°C) followed by PMSF, iv) EVs treated with proteinase K, and v) EVs samples incubated with proteinase K followed by PMSF. Immediately after treatment, EVs proteins profiles were analyzed by SDS-PAGE 12% followed by Coomassie Blue Staining.

### Dot blot

Bacterial and EV lysates were prepared according to the following procedure. Overnight cultures of *S. aureus* N315 in 10 mL of RPMI+LB were centrifuged (6,000 x *g*, 10 min, 4°C). The pellets were washed with PBS, centrifuged under the same conditions, resuspended in 1 mL of 1X RIPA buffer containing 0.1 mm zirconia beads (BioSpec Product), and shaken in an Eppendorf^®^ TherMomixer^®^ C (2,000 rpm, 30 min, 4°C). The lysates were then centrifuged (10,000 x *g*, 20 min, 4°C), and the resulting supernatant was collected for further analyses. EVs samples were diluted (v/v) with 2X RIPA buffer, and shaken in an Eppendorf^®^ TherMomixer^®^ C (2,000 rpm, 30 min, 4°C). Protein quantification was performed with Pierce^TM^ BCA Protein Assay Kits. Serial ½ dilutions of samples were deposited on nitrocellulose membranes (Hybond C extra Amersham). MG-63 and *E. coli* lysate were used as negative control (NC) for PG and lipoteichoic acids (LTA) detection, respectively. For positive control (PC), purified *S. aureus* LTA (tlrl-pslta, Invivogen) and PG (tlrl-pgns2, Invivogen) were used. For peptidoglycan detection, the upper concentration start from 10 µg for PC and 0.25 µg for SA, EVs and NC and for lipoteichoic acids from 160 ng for PC, 2 µg for SA and NC, and finally 250 ng for EVs. Membranes were incubated with 5% (w/v) of Bovine Serum Albumin (BSA) in TBS with 0.03% Tween20 (TBS-T) overnight at 4°C, followed by incubation 2 h at room temperature with either Gram-positive bacteria lipoteichoic acids monoclonal antibody (G43J, Invitrogen) diluted 1/200 in 5% BSA TBS-T or peptidoglycan monoclonal antibody (3F6B3, Invitrogen), diluted 1/1000 in 5% BSA TBS-T. Membranes were washed with TBS-T and incubated for 2 h at RT with goat anti-mouse IgG1 (γ1) (A10551, Invitrogen), horseradish peroxidase conjugate, diluted 1/2000. Membranes were washed with TBS-T, developed using Clarity Western ECL Substrate (Bio-Rad) and imaged with a ChemiDoc MP Imaging System (Bio-Rad).

### Mass spectrometry, protein identification, and protein sequence analysis

Nano-LC-ESI MS/MS experiments and analysis, including samples preparation and in-gel digestion were performed as described previously (42). The searches were performed against the predicted proteome of *S. aureus* N315 (Accession: NC_002745). The E-value threshold for peptide identification was set to 0.05 and a minimum of two peptides per replicate was required for protein identification, resulting in a false discovery rate (FDR) of < 0.15%. For each experimental condition, 3 biological replicates were investigated and the proteins were considered present if they were identified in at least 2 out of 3 replicates. Ortholog-based annotation was obtained with eggNOG-mapper (43, 44), including the assignment to clusters of orthologous groups (COG) categories, and PSORTb prediction. Lipoprotein signals were predicted with PRED-LIPO (45). Venn diagrams were obtained using Draw Venn Diagram [http://bioinformatics.psb.ugent.be/webtools/Venn/]. The mass spectrometry proteomics data can be found at https://doi.org/10.57745/OHZDZY.

### Labeling of EVs

For single membrane labelling of EVs, 15 µL of EVs sample (around 1.0 × 10^12^ particles/mL) were stained with 10 µM of Vybrant^TM^ DiI solution (V22889, Thermo Fisher Scientific) for 30 min at 37°C. For double protein- and membrane-labelling of EVs, the same amount of EVs were stained with 1 µL of biotracker^TM^ red exosome protein labeling kit (SCT269, Sigma) and 10 µM Vybrant^TM^ DiO solution (V22889, Thermo Fisher Scientific) for 30 min at 37°C. Samples were then sequentially treated with proteinase K and PMSF to remove contaminant proteins, as previously described. Labeled EVs were filtered using Amicon Ultra centrifugal filter units with 100 kDa cut-off by centrifugation (14,000 x *g*, 4 min). The flow-through was discarded and PBS was added to the retained EVs. The tubes were centrifuged again at the same speed for 3 minutes in order to wash EVs and remove unreacted dye. Centrifugation and wash step were repeated three times under the same conditions. Retained EVs were finally recovered, and particle concentrations were determined by NTA. For labelling of both RNA and membrane of EVs, 50 µL of EVs sample (around 1.0 × 10^12^ particles/mL) were stained with 10 µM Syto^TM^ RNASelect^TM^ (S32703, Thermo Fisher Scientific) and 10 µM Vybrant^TM^ DiD for 30 min at 37°C. Samples were then sequentially treated with RNase A and the RiboLock RNase inhibitor as previously used. Labelled-EV samples were subsequently purified using the SEC-based isolation protocol, concentrated, and analyzed by NTA as described above. Double-labelled EVs were analyzed by confocal microscopy; a drop of approximately 2 µL of labeled EV samples was deposited on a microscope slide with Prolong^TM^ Glass Antifade Mountant (P36980, Thermo Fisher Scientific).

### Maintenance of eukaryotic cells

Adherent fibroblast derived from human osteogenic sarcoma MG-63 cells (LGC Standards, Teddington, UK) were cultured in DMEM (D0819, Sigma) supplemented with 10% (v/v) fetal bovine serum (FBS-HI-12A, Cliniscience) and 1% (v/v) penicillin/streptomycin (P0781, Sigma). HEK-Blue^TM^ hTLR2 cells containing an inducible NF-κB/AP1 secreted alkaline phosphatase (SEAP) reporter (hkb-htlr2, InvivoGen) and the parental HEK-Blue^TM^ Null1-k cells (hkb-null1, InvivoGen) were maintained in DMEM supplemented with 10% (v/v) fetal bovine serum and 1% (v/v) penicillin/streptomycin with 100 µg/mL of normocin (ant-nr-05, Invivogen). Culture media were supplemented with HEK-Blue™ Selection (hb-sel, Invivogen), a solution containing several selection antibiotics provided by supplier and 100 μg/mL of zeocin (ant-zn-05, Invivogen) for HEK Blue^TM^ hTLR2 and parental cells, respectively. All incubations were at 37°C in 5% CO2. Trypsin/EDTA (T4049, Sigma) was used at 0.25% to release MG-63 for cells subculturing. For HEK cell line, PBS was used instead of trypsin/EDTA to preserve TLRs from degradation.

### Secreted alkaline phosphatase (SEAP) reporter assays

HEK-Blue^TM^ cells were seeded in 96-well plates at 4 × 10^4^ cells/well in a final volume of 200 µL. After 24 h, cells were stimulated with purified EVs at different doses (from 5 × 10^5^ to 5 × 10^7^ particles/well) or 10 ng/mL FSL-1 (Invivogen, tlrl-fsl) for 24 h. FSL-1 treated cells were used as a positive control. Cell culture supernatants were assayed for SEAP activity by incubating 20 μL of supernatant with 180 μL of QUANTI-Blue solution (Invivogen) at 37°C, according to the manufacturer’s instructions. SEAP activity was measured at 630 nm with the Agilent BioTek Synergy H1 microplate reader.

### Flow cytometry analysis of MG-63 cell interactions with N315 EVs

MG-63 cells were seeded in 24-well plates at density of 3.0 × 10^4^ cells per cm² in a final volume of 1 mL and cultured for 48 h. The medium was then changed and supplemented, or not, with 80 µM dynasore (324410, Sigma), 10 µM chlorpromazine (31679, Sigma), 20 µM nystatin (N6261, Sigma,) or 5 µM cytochalasin (C8273, Sigma), one hour prior to exposure to varying amounts of DiI-labeled EVs (from 2.5 to 40 µg) for incubation times ranging from 0.5 to 17 h. Each well was washed twice with PBS, and cell pellet were recovered after trypsinization and centrifugation. Cell pellets were fixed with 4% PFA (20 min, RT in the dark), and washed twice with PBS. Samples were analyzed on a BD LSRFortessa X-20 flow cytometer equipped with five lasers (UV 355 nm, Violet 405 nm, Blue 488 nm, Yellow-Green 561 nm, and Red 640 nm). Total cell counts were determined by gating on SSC-A versus FSC-A plots, with cellular debris excluded from intact cells by proper gating. For DiI, excitation was performed using the 561 nm laser with a 586/15 emission filter. DiI fluorescence intensity (indicative of EV internalization) was evaluated from histogram plots of PE versus count, with PE-A set on a logarithmic scale and count on a linear scale. MG-63 cells incubated with PBS only or with unlabeled EVs were used as an unstained control for gate setting. Data were analyzed with FSC Express software and expressed as an internalization factor, calculated as the median fluorescence multiplied by the percentage of marker-positive events.

### Confocal microscopy analysis of MG-63 cell interactions with N315 EVs

MG-63 cells were seeded in wells of Nunc^TM^ Lab-Tek II chamber slide^TM^ system 8 (ThermoFisher scientific) at densities of 3.0 × 10^4^ cells per cm². After 48 h, the medium was changed, and supplemented, or not, with dynasore or chlorpromazine, one hour prior to exposure to 20 µg of labeled EVs (MOI 1000 for endosomal experiments) for incubation periods of 3 h or 22 h. Cells were washed twice with PBS, fixed with 4% PFA, and washed twice again with PBS. Cell membranes were stained successively with Phalloidin-iFluor 488 Reagent (ab176753, Abcam) (1/1000 dilution, 1h, RT) and then with DAPI included in mounting medium (VECTASHIELD® PLUS Antifade Mounting Medium with DAPI, Eurobio Scientific, H-2000-10) with washing step in between. Slides were stored at 4°C until analysis, and imaged with a confocal microscope (LSM880, Carl Zeiss, France) using appropriate lasers and objectives : 20X/0.8 NA objective or 63X/1.4 NA oil objective.

### RT-qPCR analysis of MG-63 cell interactions with N315 EVs

MG-63 cells were seeded in 24-well plates at density of 3.0 × 10^4^ cells per cm² in 1 mL of medium, and cultured for 48 h. The medium was changed, and supplemented, or not, with 80 µM dynasore, one hour prior to exposure to PBS (control for EV treatment), 0.2% dimethyl sulfoxide (DMSO), 20 µg of EVs in PBS, or 20 µg of EV in PBS with 0.2% DMSO (final concentration) (control for EV treatment in the presence of dynasore). After 3 h of incubation, the culture medium was discarded, and cells were washed twice with PBS. Total RNA was extracted using the RNeasy Mini kit (Qiagen, 74106), and treated with DNase I (AMPD1-1KT, Sigma-Aldrich,) according to manufacturer’s instructions. One µg of DNA-free RNA was reverse transcribed by qScript cDNA Synthesis kit (Quantabio) for a final volume of 20 µL according to the manufacturer’s instructions, and qPCR was carried out qPCR was carried out in a 15 μL volume containing 15 ng cDNA, specific primers (300 nM), and 8 μL IQTM SYBR Green Supermix (Bio-Rad). Primers used are listed in Table S2. Reactions were run on a CFX96 real-time system (Bio-Rad, France) using the following cycling parameters: DNA polymerase activation and DNA denaturation 95°C for 3 min, 40 cycles of denaturation at 94°C for 10 s, and extension at 60°C for 40 s. Melting curve analysis was included to check the amplification of single PCR products. All primers used were described in supplementary data **(Table S2)**. RT-qPCR was conducted using three biological replicates, and three independent technical replicates (n = 9) for each cDNA sample. The amount of mRNA was normalized by using the geometrical mean value of three reference genes, *GAPDH*, *PGK1*, and *PPIA*. The online tool RefFINDER (46) was used to identify these genes as the most stably expressed genes in the experimental conditions tested from a panel of five candidate genes. Gene expression values were calculated using the 2^−ΔΔCT^ method relative to control conditions (PBS for EV treatment and EVs in PBS-DMSO for EV treatment in the presence of dynasore) and expressed as mean ± SD. RT-qPCR data can be found at https://doi.org/10.57745/AZR96A.

### NF-κB pathway array

MG-63 cells were seeded in 24-well plates at density of 3.0 × 10^4^ cells per cm² in 1 mL of medium, and cultured for 48 h. After change of medium, PBS or N315 EVs (20 µg/well) were added and incubated for 24h. Then, the culture medium was discarded, and cells were washed twice with PBS. One hundred µL of lysis buffer from Human Proteome profiler NF-κB Pathway Array Kit (ARY029, R&D Systems, Bio-techne) were added per well and kept 5 minutes on ice. Afterwards, 6 wells were pooled per conditions and placed at 4°C for 30 minutes with strong agitation. Samples were centrifuged (14,000 x *g*, 5 min, 4°C), and supernatants were recovered. Proteins quantification was performed using Pierce^TM^ BCA Protein Assay kit (A55864, Thermo Scientific). The rest of the experiment were conducted according to the manufacturer’s protocols using 200 µg of proteins derived from the lysate of MG-63 cells stimulated for 24 hours with PBS or N315-derived EVs. Membranes were developed using Clarity Western ECL Substrate (Bio-Rad) and imaged with a ChemiDoc MP Imaging System (Bio-Rad). Integrated pixel density for each spot was measured using Fiji (ImageJ 1.54p, Java 1.8.0_452 (64bit)) (47).

### Image analysis

Image analysis was performed using Fiji (ImageJ 1.54p, Java 1.8.0_452 (64bit)) (47). Confocal microscopy images were converted to 8-bit format, adjusted for brightness and contrast, before being smoothed using a Gaussian blur filter (sigma radius = 1). For endosomes quantification part, resizing of all images was performed without interpolation to allow comparison. ROIs were defined manually on the DAPI channel images, which mark both cell membranes and nuclei. The images from the relevant channels (RNA, endosomes, and EVs-membranes) were opened separately. A threshold was applied using manual adjustment in the same way for consistency. Afterwards, images were converted to binary format and every pixel with two or three labels was counted inside each ROI and reported on the total number of pixels with each of the individual label. For instance, pixels for EVs with double-labelling were reported on all the corresponding pixels:

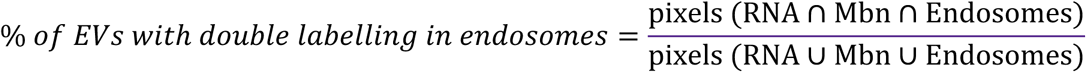

### Statistical analysis

Data analysis was performed using GraphPad PRISM 10 for macOS (version 10.6.0). All data are presented as mean ± standard deviation (SD). Normality tests were performed using Shapiro–Wilk tests, while F tests were used to compare variances. Depending on data distribution, unpaired t-test, Welch’s t test, or Mann-Whitney test was used to analyze the statistical difference between groups of two. Statistical analyses for more than two groups were performed with one-way ANOVA followed by Tukey’s or Dunnett’s multiple comparisons test. Gray-value data for line plot were obtained with Fiji and smoothed with GraphPad (6 neighbors on each size and 0th order of the smoothing polynomial).

## RESULTS

### *S. aureus* N315 strain produces EVs *in vitro*

The ability of *S. aureus* strain N315 to produce EVs was assessed in RPMI, a medium physiologically relevant for mammalian cell culture, supplemented with 10% LB to support bacterial growth (Fig. S1). Particles concentration in the culture medium was quantified prior to bacterial inoculation and after 16 hours of growth to monitor EV production over time. As shown in Figure 1A, N315 growth induced a > 30-fold increase in particle concentration, rising from 5 × 10^7^ to 1.5 × 10^9^ particles per mL of culture. Moreover, the size distribution of the particles detected post-incubation differed from that of those initially present, further supporting the active release of EVs by strain N315 during growth in RPMI+LB (Fig. 1B, left and middle panels). Culture supernatants were then subjected to EV isolation procedures using size exclusion chromatography. The fractions collected were carefully selected to ensure the best compromise between EV yield and sample purity, minimizing protein contamination (Fig. S2) and the pool of interest fractions was characterized by NTA and electronic microscopy. NTA revealed particles within the 90-110 nm size range, with concentrations ranging from 2.6 to 3.4 × 10^8^ particles per mL of culture (Fig. 1A and B). Electron microscopy confirmed the presence of nanosized, membrane-bound vesicles with a typical spherical shape (Fig. 1C). These findings showed the capacity of strain N315 to produce a large amount of EVs in RPMI+LB.

**FIG. 1.**
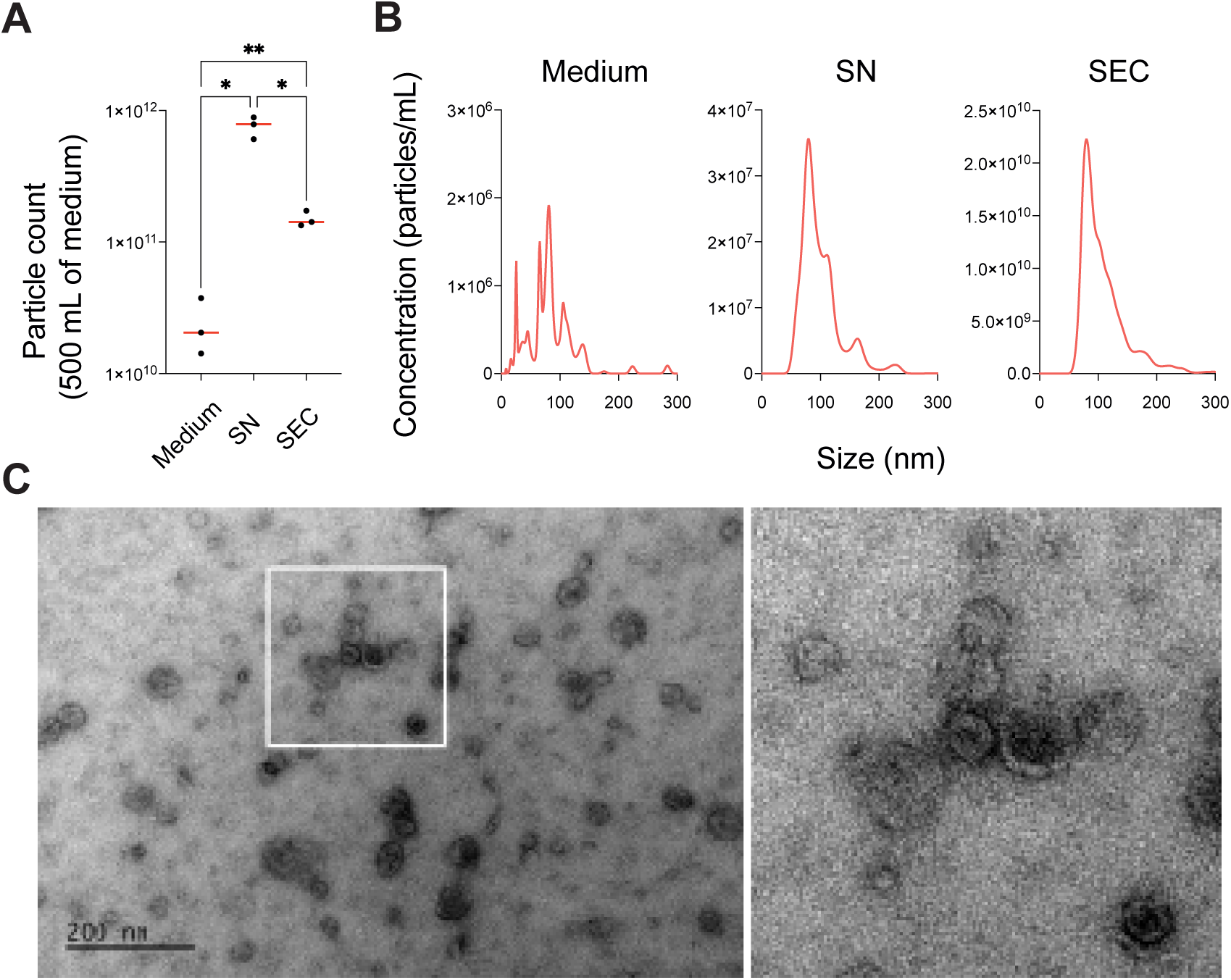
Methicillin-resistant human *S. aureus* N315 strain releases EVs *in vitro*. (A) Quantification of particles in 0.5 L of RPMI+LB medium before N315 inoculation (Medium), filtered N315 culture supernatant after 16 h of growth (SN) and following EV isolation by size exclusion chromatography (SEC). Data and graphs were obtained by NTA from three independent biological replicates. Significance was assessed by one-way ANOVA with Dunnett’s multiple comparisons test: *p* <0.05 (*); *p* < 0.01 (**). (B) Size distribution profiles of particles detected in RPMI+LB before N315 inoculation (Medium), in the filtered N315 culture supernatant after 16 h of growth (SN) and following EV isolation (SEC). Graphs are representative of 3 independent biological replicates revealed by NTA. (C) Representative electron microscopy image of N315 EVs isolated via SEC. Scale bar: 100 nm.

### *S. aureus* N315 EVs contain a diverse, selectively enriched and protected molecular cargo

EVs derived from N315 were found to contain a diverse molecular cargo including PG, LTA, DNA, RNA, and proteins (Fig. 2A). On average, 10^10^ N315 EVs contained 7 μg of protein, 70 ng of RNA and 40 ng of DNA. Proteomic analysis by LC-MS/MS identified 91 proteins associated with EVs, predominantly of cytoplasmic origin such Eno, and lipoproteins (Fig. 2B to 2E). Among them, twenty-six were not detected in the whole-cell proteome, indicating their selective enrichment within EVs. To confirm the packaging of RNA and proteins into EVs, both nuclease and protease protection assays were performed (Fig. 2F). The resulting profiles of nuclease protection revealed that EVs contained abundant intravesicular RNA, predominantly small RNAs (< 200 nt), with a major peak around 100 nt. For the protease protection assay, EVs were exposed to proteinase K under conditions that preserved or not their integrity (*i.e*., +/− RIPA). The protection from degradation of proteins only when EV integrity was maintained indicated the intravesicular protein localization and the lipid bilayer protective effect. The intravesicular localization of RNAs and proteins was further confirmed by confocal microscopy (Fig. 2G). EVs were treated to remove external proteins and RNAs, then double-labeled using a membrane-specific dye (*i.e.*, DiO or DiD) and either a lipid-permeable RNA dye (staining (*i.e.*, Syto RNA Select) or protein-specific dye (*i.e.*, BioTracker™ Far Red Exosome Protein Labeling Kit). Confocal microscopy revealed co-localization of both dyes within the EVs, supporting the presence of intravesicular RNA and protein protected by the lipid membrane. Finally, to illustrate selective protein packaging, the protein Eno, previously identified as associated with N315 EVs by LC-MS/MS, was tagged with sfGFP and its localization within EVs was assessed by confocal microscopy. As observed for RNA and total protein staining, colocalization of the sfGFP and membrane signals confirmed the incorporation of Eno into EVs. Our results highlighted the diverse molecular cargo of N315-derived EVs and the selective intravesicular localization of proteins and RNAs.

**FIG. 2.**
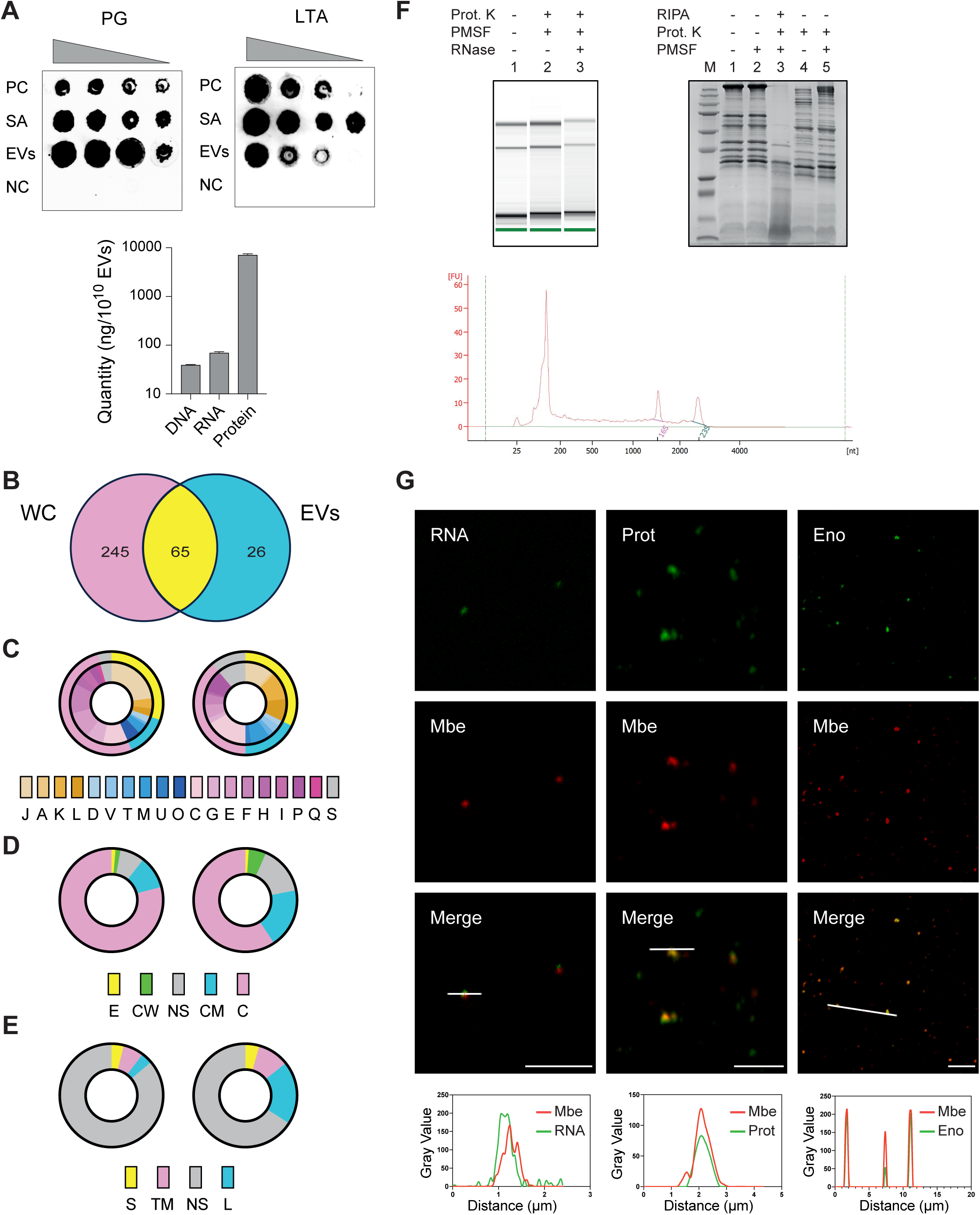
N315 EVs contain a diverse, selectively enriched, and protected molecular cargo. (A) Detection and quantification of pathogen-associated molecular patterns (PAMPs) and biomolecules associated with N315 EVs. Upper panel: dot blot detection of peptidoglycan (PG) and lipoteichoic acid (LTA). PC, positive control (commercial LTA/PG); SA, *S. aureus* N315 whole-cell lysate; EVs, purified N315 EVs; NC, negative control (*E. coli* lysate for LTA and MG-63 lysate for PG). Serial ½ dilutions were loaded. Lower panel: quantification of DNA, RNA and proteins. Data represent mean ± SD of three biological replicates. (B-E) Comparative proteomic analysis of *S. aureus* whole-cells (WC, right panels) and EVs (EVs, left panels). (B) Venn diagram showing the number of proteins identified in WC, EVs and their overlapp. (C) Functional classification of WC and EV proteins based on clusters of orthologous groups (COG) categories, predicted using eggNOG-mapper. J, translation, ribosomal structure, and biogenesis; A, RNA processing and modification; K, transcription; L, replication, recombination, and repair; D, Cell cycle control, cell division, chromosome partitioning; V, defense mechanisms; T, signal transduction mechanisms; M, cell wall/membrane/envelope biogenesis; U, intracellular trafficking, secretion and vesicular transport; O, post-translational modification, protein turnover and chaperones; C, energy production, and conversion; G, carbohydrate transport and metabolism; E, amino acid transport and metabolism; F, nucleotide transport and metabolism; H, coenzyme transport and metabolism; I, lipid transport and metabolism; P, inorganic ion transport and metabolism; Q, secondary metabolites biosynthesis, transport, and catabolism; S, function unknown. (D) Predicted subcellular localization of proteins using PSORTb. E, extracellular; CW, cell wall; NS, no signals found; CM, cytoplasmic membrane; C, cytoplasmic. (E) Lipoprotein prediction with PRED-LIPO. S, surface; TM, transmembrane; NS, no signals found; C, cytoplasmic; L, lipoproteins. (F) Nuclease and proteinase K protection assays. Upper left panel: RNA profiles from Bioanalyzer prokaryote total RNA Pico Assay: line 1, untreated EVs; lane 2, EVs treated with proteinase K + PMSF; lane 3, EVs treated with proteinase K + PMSF + RNaseA + RiboLock® inhibitor. Lower panel: Electropeherogram of lane 3 showing predominant small RNAs (< 200 nt). FU, fluorescence units; nt, nucleotide length. Upper right panel: SDS-PAGE analysis of EV samples untreated or treated with phenylmethylsulfonyl fluoride (PMSF), proteinase K (Prot. K), and/or RIPA. M, protein marker. (G) Confocal microscopy visualization of intravesicular RNA and proteins. RNA was visualized by treating EVs with RNaseA followed by staining with SytoRNA Select (RNA dye) and DiD (lipid dye). Proteins were visualized by treating EVs with proteinase K followed by staining with BioTracker Red Exosome Protein Labeling kit (protein dye) and DiO (lipid dye). EVs from the eno-sfGFP reporter strain were labeled with DiD and imaged for GFP-tagged enolase. Line graphs show fluorescence intensity profiles across individual EVs. White arrows indicate co-localized signals within the vesicle lumen.

### *S. aureus* N315 EVs are recognized by host cells via various PRRs

The molecular characterization of N315 EVs revealed the presence of numerous components able to interact with host cells, notably various PAMPs such as LTA, PGN, lipoproteins and nucleic acids, supporting their potential role in bacteria-host interactions. To gain further insight into how *S. aureus* EVs signal and interact with host cells during infection, the ability of N315 EVs to activate human PRRs was assessed *in vitro* using MG-63 cells. Specifically, we analyzed the expression of several genes coding for Toll-like receptors (TLRs) and nucleotide oligomerization domain (NOD)-like receptors (NLRs) in the presence or absence of N315 EVs (Fig. 3). Compared to unstimulated control, EVs significantly increased the expression of several *TLR* genes, including those coding for the extracellular receptors TLR1, TLR2, TLR4, and TLR6, as well as the endosomal receptors TLR3, TLR4 and TLR7. In contrast, the expression levels of *TLR5* and *TLR9* genes remained unchanged. In addition, EVs increased the expression of the cytosolic receptor *NOD2*, while *NOD1* expression was unaltered. Notably, for *TLR2*, the EV-induced gene upregulation was confirmed at the protein level (Fig. 4A and B). To further demonstrate a direct interaction between EVs and PRRs, we used a reporter cell line, the HEK-Blue^TM^ hTLR2 cells, which were designed for studying the stimulation of the human TLR2 receptor by agonists or antagonists by monitoring the activation of NF-kB transcriptional regulator. Stimulation with increasing concentrations of EVs, resulted in a dose-dependent increase of SEAP reporter activity, showing the direct interaction between EVs and TLR2 (Fig. 4C). Together, these findings showed the ability of N315-EVs to activate various PRRs located at the cell surface, within endosomes, and in the cytoplasm of cells, highlighting the potential EV role in modulating innate immune responses.

**FIG. 3.**
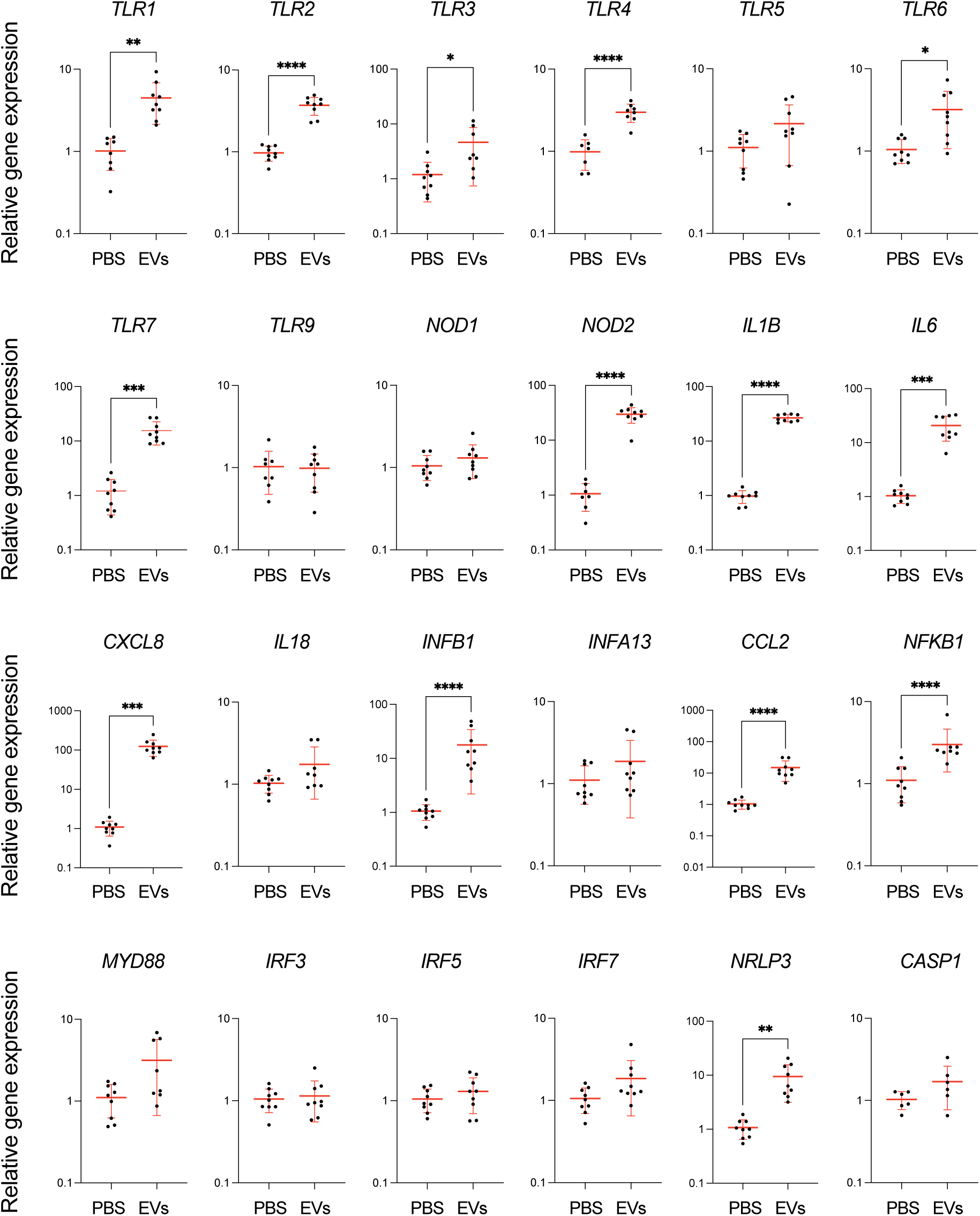
N315 EVs modulate the expression of immune-related genes in MG-63 cells. Relative fold change in mRNA expression of PPRs (TLR1, TLR2, TLR3, TLR4, TLR5, TLR6, TLR7, TLR9, NOD1, NOD2), cytokines and chemokines (IL1B, IL6, CXCL8, IL18, INFB1, INFA13, CCL2), and components of signaling pathways (NFKB1, MYD88, IRF3, IRF5, IRF7, NRLP3, CASP1) in MG-63 cells after 3 h stimulation with N315 EVs *versus* PBS-treated controls. Relative gene expression was calculated by the 2^−ΔΔCt^ method, using PBS-treated cells as control and *GAPDH*, *PGK1*, and *PPIA* genes for normalization. Data are presented as mean ± SD of biological triplicates analyzed in triplicate (n = 9). Statistical analyses were performed using unpaired t-test or Welch’s t-test when required: *p* <0.05 (*); *p* <0.01 (**); *p* <0.001 (***); *p* <0.0001 (****).

**FIG. 4.**
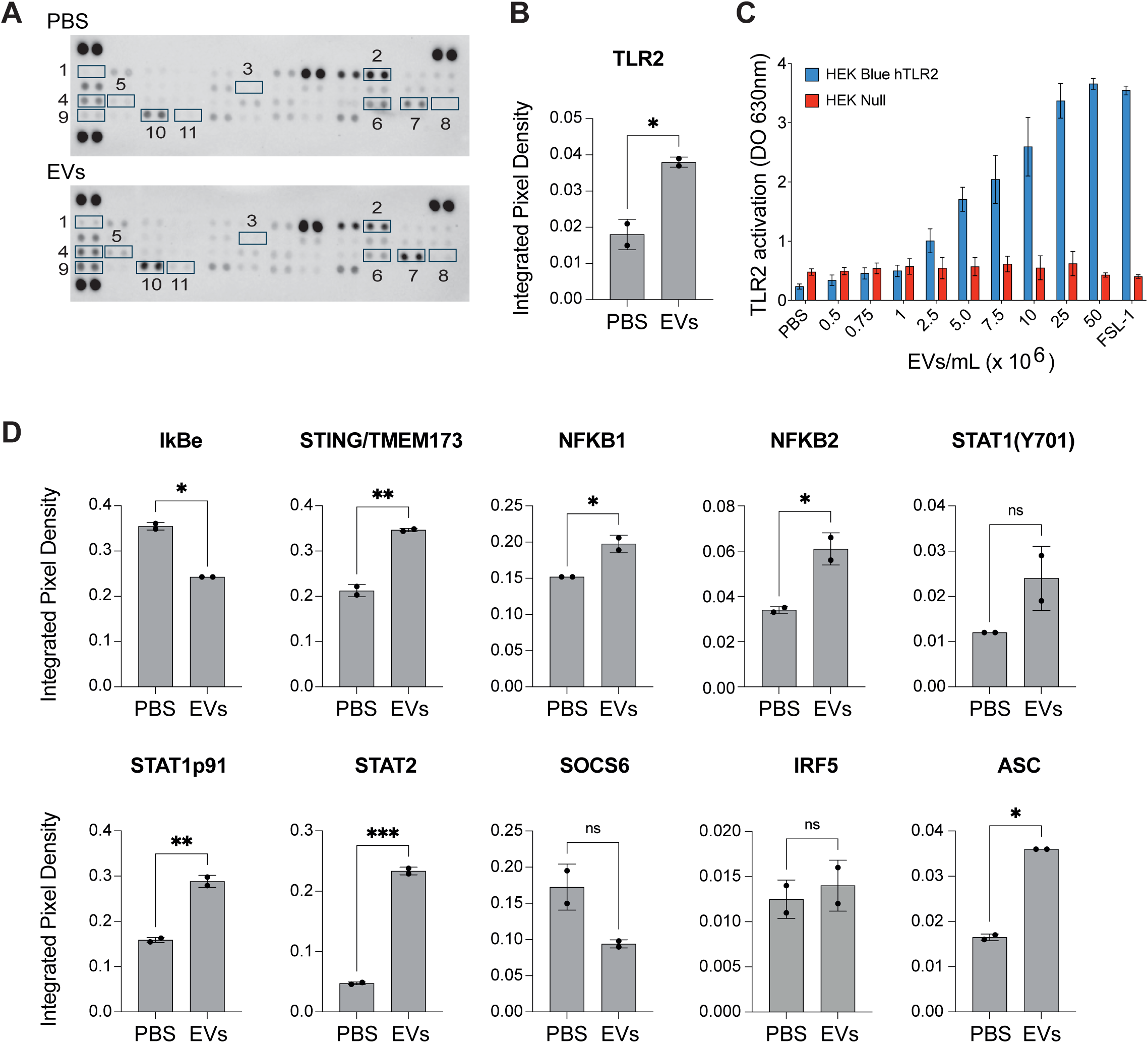
N315 EVs activate the NF-kB and JAK/STAT signaling pathways in MG-63 cells. (A) Membrane-based antibody array analysis of MG-63 cell lysates (24 h post-EV exposure). Spots correspond to: 1, ASC; 2, IκBε; 3, IRF5; 4, NF-κB1; 5, NF-κB2; 6, SOCS6; 7, STAT1-p91; 8, STAT1-pY701; 9, STAT2; 10, STING/TMEM173; 11, TLR2. Upper panel: PBS control; Lower panel: 20 µg N315 EVs. (B) Quantification of integrated pixel density for TLR2 from (A). Each antibody target was spotted in duplicate. Data are presented as mean ± SD of replicates. Statistical analyses were performed using unpaired t test or Welch’s t-test when required: *p* <0.05 (*); *p* < 0.01(**); *p* < 0.001(***). (C) Activation of TLR2 by N315 EVs using HEK-Blue^TM^ hTLR2 reporter cells. SEAP activity was measured after 24 h of stimulation with increasing EV concentrations. FSL-1 (10 ng/mL) was a positive control; HEK-Blue^TM^ Null1 cells were used as a negative control. Data represent mean ± SD of three biological replicates. (D) Quantification of integrated pixel densities for all other targets shown in (A). Data are presented as mean ± SD of replicates. Statistical analyses were performed using unpaired t-test or Welch’s t-test when required: *p* < 0.05 (*); *p* < 0.01 (**); *p* < 0.001 (***).

### *S. aureus* N315 EVs induce inflammation via the NF-κB and STAT/JAK signaling pathways

Activation of PRRs typically triggers downstream signaling cascades that result in the production of many cytokines and chemokines. To confirm the immunomodulatory effect of N315 EVs following PRRs activation, we first assessed their ability to induce cytokine expression. Exposure of MG-63 cells to N315-EVs resulted in a significant upregulation of *IL1B*, *IL6*, *CXCL8, CCL2*, and *INFB1* genes primarily regulated by the NF-κB signaling pathway. The activation of the NF-κB signaling pathway by N315 EVs was further supported by the increased expression of *NFKB1* gene (Fig. 3), together with the decrease in the abundance at the proteomic level of IκBε (NFkB inhibitor) and the concomitant increase of STING, NFKB1, and NFKB2, 24 h after EV exposure (Fig. 4D). Beyond NF-κB activation, our data suggest that N315-EVs may also signal throughout the JAK/STAT signaling pathway. Indeed, proteomic analysis revealed a strong upregulation of STAT1 (Y701 and p91 isoforms) and STAT2, two key transcription factors of the JAK/STAT pathway (Fig. 4D). In line with this, we observed a tendency toward downregulation of SOCS6 protein, a member of the suppressor of cytokine signaling (SOCS) family, involved in the negative regulation of this pathway. Finally, in contrast, the unchanged expression of *IRF3*, *IRF5*, *IRF7* and *INFA13* (Fig. 3) and production of IRF5 (Fig. 4D) in the presence of EVs suggested that the IRF pathways were not involved in the cell response to EV exposure. Altogether, our findings highlight the capacity of N315 EVs to modulate host immune responses primarily through NF-κB and JAK/STAT pathways, while bypassing IRF-mediated signaling, triggering a persistent pro-inflammatory profile.

### *S. aureus* N315 EVs induce inflammasome

EV exposure induced the expression of several proteins associated with cell death pathways. A slight but non-significant upregulation of death receptors like Fas, TNFR1, TRAILR1, TRAILR2, DR3, and NGFR proteins was observed at 24 h post-incubation, all of which are involved in apoptosis (Fig. S3). Moreover, we observed the upregulation of *NLRP3* and *IL1B* gene expression at 3 h post-incubation suggesting the impact of N315-EVs on inflammasome induction (Fig. 3). NLRP3 forms a complex with ASC and CASP1 that activates IL1β and IL18 reportedly involved in inflammation and pyroptosis. Accordingly, we found the upregulation of *IL1B* expression at 3 h post-incubation (Fig. 3) and ASC abundance at 24 post-incubation (Fig. 4D).

### *S. aureus* N315 EVs are internalized in MG-63 cells through dynamin-dependent endocytosis

The ability of N315 EVs to activate intracellular receptors in a non-phagocytic human cell line raised questions about their uptake. To investigate this, labeled EVs were incubated with MG-63 cells stained with DAPI for cell nucleus (blue) and with Phalloidin for actin filaments (green) and analyzed by confocal microscopy (Fig. 5A). After 3 h of interaction with EVs, a colocalization of EV and cell signals was observed, suggesting EV internalization. A 3D reconstruction further confirmed the intracellular localization of EVs, notably in the perinuclear region. When compared to 3 h of exposure, we noted a clear increase in the signal after 22 hours, suggesting an increase of EV internalization over time. Flow cytometry analysis supported these findings, showing that the uptake of membrane-labeled EVs by MG-63 cells increased in a time- and dose-dependent manner (Fig. 5B and C). To gain insight into the mechanisms of internalization, MG-63 cells were pretreated with specific inhibitors of internalization pathways prior to incubation with EVs: Chlorpromazine (clathrin-mediated endocytosis), dynasore (dynamin-mediated endocytosis), nystatin (caveolae/lipid raft-mediated endocytosis), and cytochalasin D (phagocytosis). It should be noted that we also tried using methyl-b-cyclodextrin, but it proved toxic to our cells at the concentrations used (data not shown). Among these, only dynasore significantly decreased EV uptake (Fig. 5D and E), indicating that N315 EVs are primarily internalized via a dynamin-dependent endocytosis pathway.

**FIG. 5.**
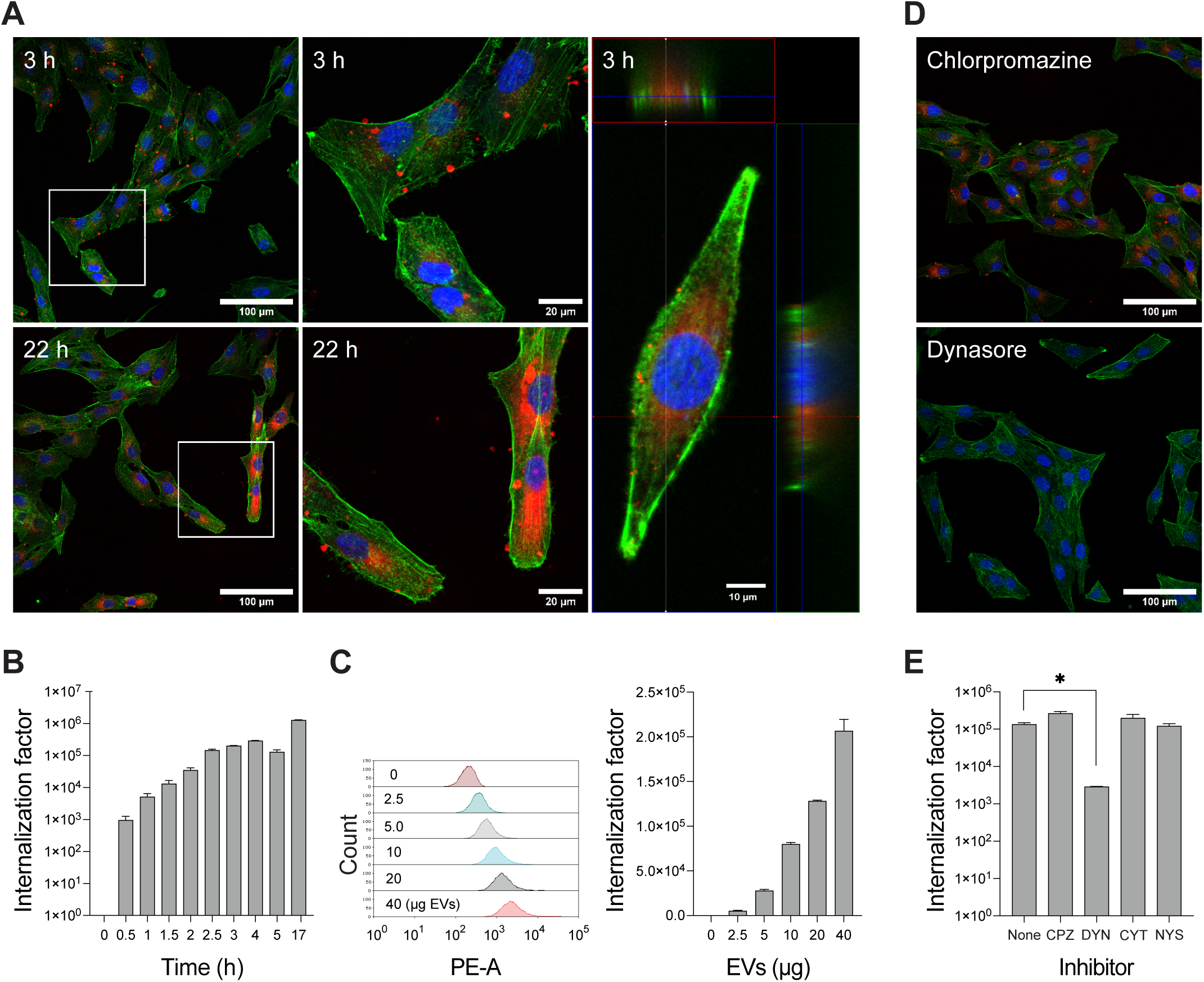
N315 EVs are internalized by MG-63 cells in a time- and dose-dependent manner by dynamin-mediated endocytosis. (A) Confocal microscopy visualization of MG-63 cells stained with Hoescht (nuclei, blue) and Phalloidin (actin filaments, green) after incubation with 20 µg DiI-labelled EVs (membrane, red) for 3 h and 22 h. Right panel: 3D reconstruction showing the intracellular distribution of EVs. (B) Flow cytometry analysis of MG-63 cells incubated with DiI-labelled EVs at different incubation time. (C) Flow cytometry analysis of MG-63 cells after 3 h of incubation with increasing quantities of DiI-labelled EVs. (D) Confocal microscopy visualization of MG-63 cells stained with Hoescht (nuclei, blue) and Phalloidin (actin filaments, green) after incubation with 20 µg DiI-labelled EVs (membrane, red) for 3 h in the presence of chlorpromazine or dynasore. (E) Flow cytometry analysis of MG-63 cells after 3 h of incubation with 20 µg DiI-labelled EVs and with or without pre-treatment with chlorpromazine (CPZ), dynasore (DYN), cytochalasin (CYT) or nystatin (NYS). Data are presented as mean ± SD of three technical replicates for time-course experiments and two technical replicates for EV dose-response and inhibitor assays.

### *S. aureus* N315 EVs release their cargo within late endosome following internalization

To investigate the intracellular fate of EVs and their cargo, double-labeled EVs, *i.e.*, membrane-stained and labeled for either RNAs or proteins, were incubated with MG-63 cells pre-labeled for nuclei and membrane and analyzed by confocal microscopy (Fig. 6). After 3 hours of interaction, signal colocalization between the two markers (membrane/RNAs or membrane/proteins) was detected in MG-63 cells, with a characteristic perinuclear distribution (Fig. 6A). The observed colocalization of EV signals supports the presence of intact EVs within MG-63 cells. To confirm the presence of EVs within endosomal compartments in MG-63 cells, similar confocal imaging experiments were carried out using late endosome labeling in cells incubated with double-labeled EVs, as previously described (Fig. 6B). A stable colocalization of fluorescence signals corresponding to EV membrane and RNA cargo was observed within cells during the first three hours of co-incubation, confirming the structural integrity of EVs. However, at six hours post-incubation, the colocalization of these signals significantly decreased, suggesting a dissociation of the EV membrane from its RNA cargo (Fig. 6C, left panel). At 1 and 3 hours, both EV membrane and RNA signals, as well as their overlap, colocalized with late endosome markers, confirming the presence of intact EVs within late endosomes (Fig. 6C, middle and right panels), as previously suggested by their perinuclear localization. By six hours, a marked reduction in the colocalization of EV membrane and RNA signals with late endosomes was observed, whether analyzed separately or together. Colocalization experiments suggested that intact EVs were present within late endosomes up to 3 hours post-incubation and that their RNA cargo was subsequently released within these compartments at later time points.

**FIG. 6.**
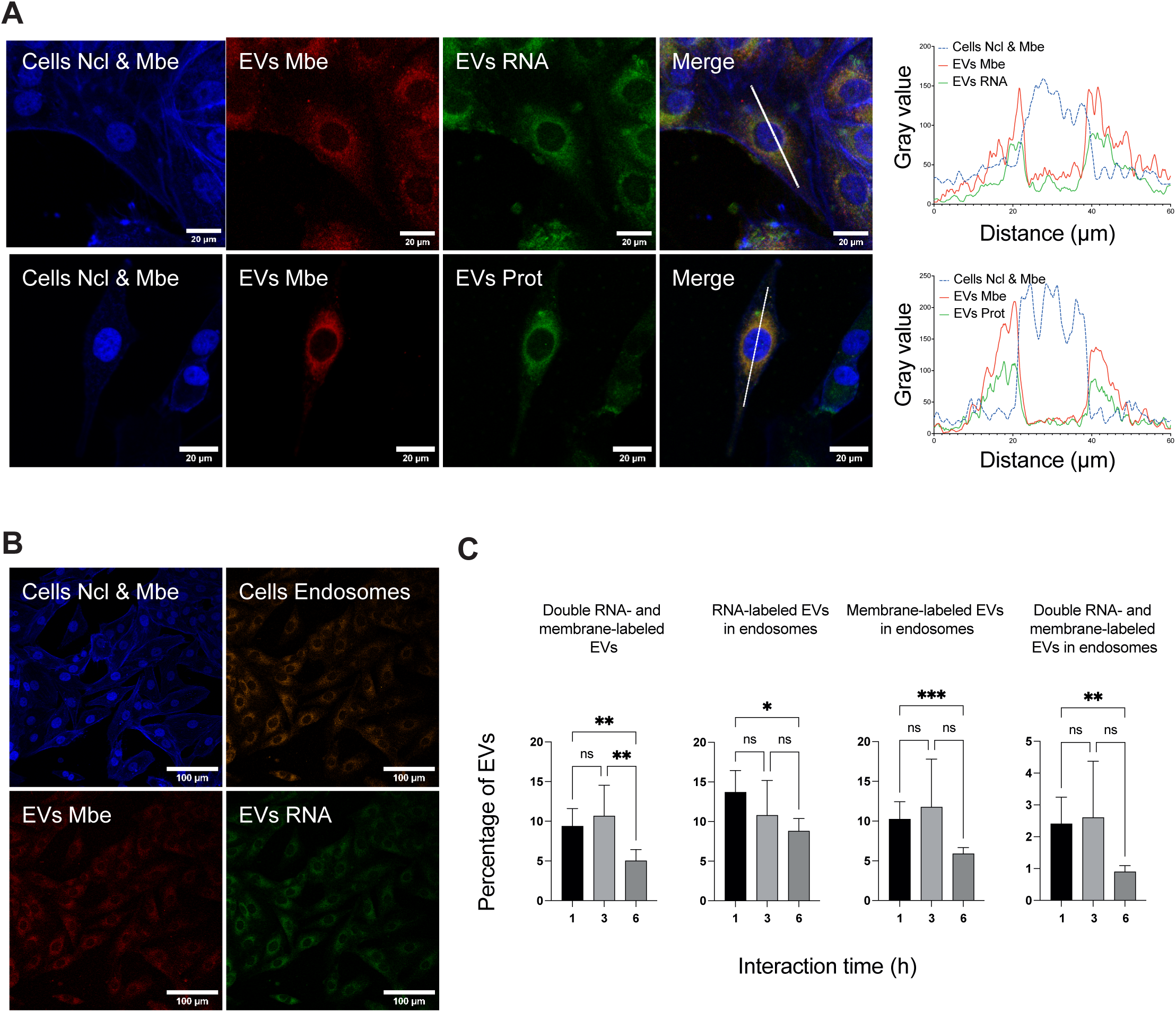
N315 EVs traffic through the endosomal-pathway and release their cargo into late endosomes. (A) Confocal microscopy visualization of MG-63 cells incubated with 20 µg double-labelled EVs for 3h. Cells were stained with DAPI, which marks both cell membranes and nuclei. EVs were stained with either DiD (EVs Mbe) and SytoRNA Select (EVs RNA) or DiO (EVs Mbe) and Biotracker (EVs Prot). Line graphs show gray value profiles along the line across the extracellular vesicles in the panel associated. (B) Confocal microscopy visualization of MG-63 cells incubated with 20 µg double-labelled EVs for 3h. Cell nuclei were stained using DAPI (blue) as previously used and late endosomes were stained with CellLights Late Endosomes-RFP (orange). EVs were stained with DiD (EVs Mbe) and SytoRNA Select (EVs RNA). (C) Quantification of EV cargo dissociation over time. Percentage of EVs exhibiting co-localization between membrane (DiD/DiO) and RNA/protein cargo within late endosomes at 1, 3, and 6 h post-incubation. Calculated as: % co-localized EVs = (Pixels (Membrane ∩ Cargo ∩ Late Endosome)) / (Pixels (Membrane ∪ Cargo ∪ Late Endosome)) × 100. Data from two biological replicates, with ≥150 cells analyzed per condition; duplicates per replicate. Error bars: mean ± SD. Statistical analyses were performed using one-way ANOVA with Tukey’s multiple comparisons test: *p* < 0.05 (*); *p* < 0.01 (**); *p* < 0.001 (***); *p* < 0.0001 (****).

### Inhibition of *S. aureus* N315 EV uptake alters their interaction with host cells

To assess the potential regulatory effects of EVs once internalized, cells were incubated with EVs in the absence and presence of dynasore, and the expression of genes previously shown to be modulated by EVs was measured by RT-qPCR (Fig. 7). Following dynasore treatment, the expression of *TLR2*, *CXCL8*, *INFB1* was upregulated, whereas *TLR3*, *TRL7*, and *IL6* were significantly downregulated. In addition, *MYD88*, *IRF3*, and *IRF7* expression were also reduced in the presence of dynasore. By contrast, the expression of other EV-dependent genes, such as *TLR1*, *NOD2*, and *NFKB1*, remained unaltered. These results show that EV internalization through the dynamin-dependent endocytosis pathway is a required step to drive the modulation of some genes involved in TLR signaling and cytokine production and that multiple entry pathways may coexist.

**FIG. 7.**
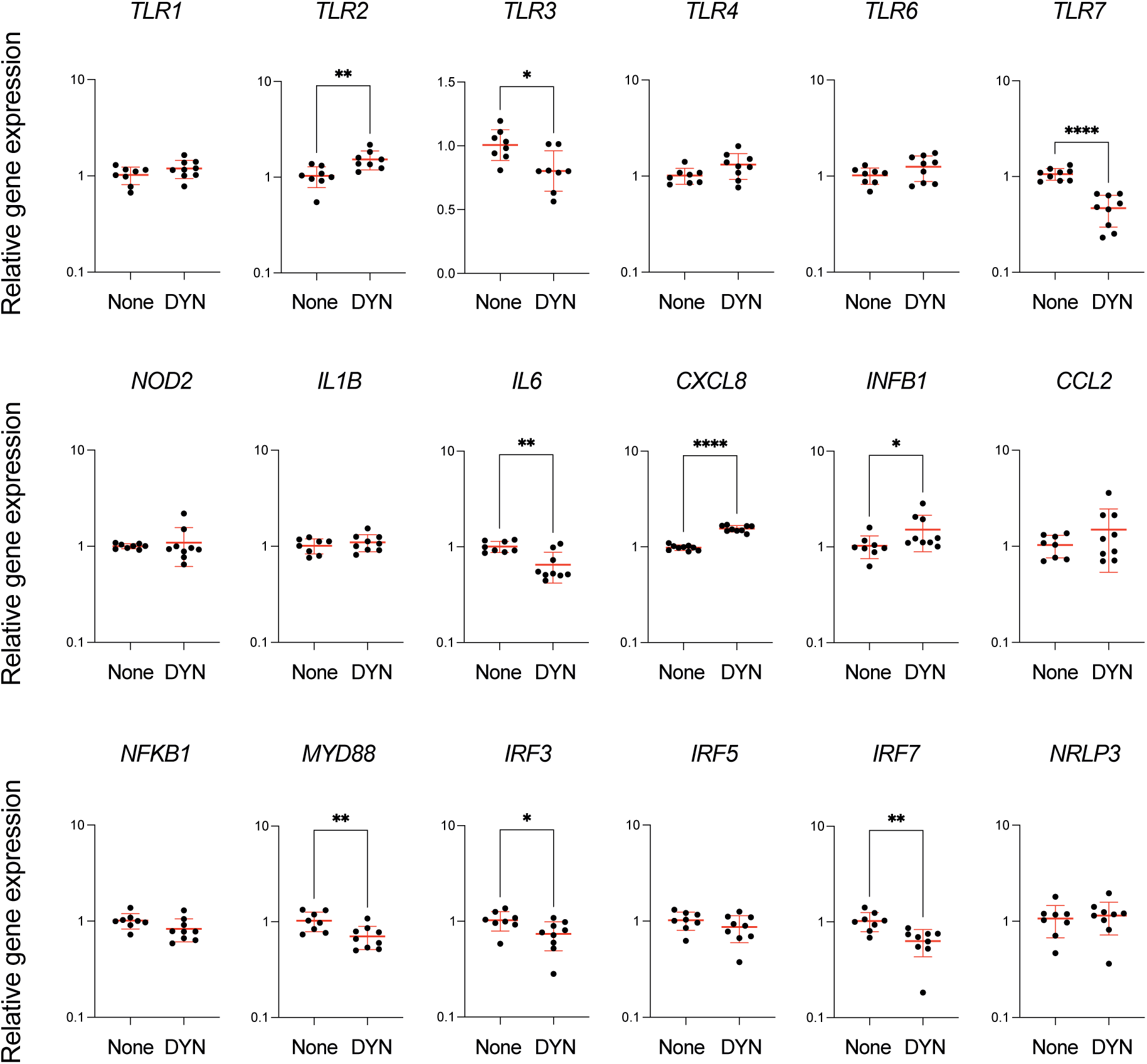
Inhibition of EV internalization alters specific immune gene expression in MG-63 cells. Relative fold change in mRNA expression of key immune genes in MG-63 cells treated with N315 EVs alone (None) or with EVs + dynasore (80 µM, DYN). Relative gene expression was calculated by the 2^−ΔΔCt^ method, using EV-treated cells as control and *GAPDH*, *PGK1*, and *PPIA* genes for normalization. Data are presented as mean ± SD of biological triplicates analyzed in triplicate (n = 9). Statistical analyses were performed using unpaired t-test or Welch’s t-test when required: *p* <0.05(*); *p* <0.01 (**); *p* <0.001 (***); *p* <0.0001 (****).

## DISCUSSION

The production and secretion of EVs by *S. aureus* are strongly influenced by growth conditions (*e.g*., growth phase, culture medium), environmental stresses (*e.g.*, H2O2, osmotic stress, antibiotics), and strain-specific factors (*e.g.*, PSM level) (48–50, 42, 51, 30). These factors not only affect the quantity of EVs produced but also shape their molecular cargo (23, 42, 49). Although some of the stress conditions tested *in vitro* may mimic those encountered by the bacterium during infection (*e.g.*, iron limitation, H_2_O_2_), most studies rely on rich laboratory media for EVs production. In this study, we evaluated the ability of *S. aureus* strain N315, a methicillin-resistant (MRSA) human isolate, to produce and secrete EVs. N315 was cultured in RPMI, an eukaryotic cell culture growth medium, supplemented with 10% LB to sustain bacterial growth. We selected these conditions because RPMI optimally promotes immune cell activation, better approximating *in vivo* responses (52), and because iron-depleted RPMI medium best recapitulates *S. aureus* infection-like conditions (53). N315 successfully produced EVs under these conditions. At this stage, however, direct comparison of EVs yields between N315 in this condition and previously reported strains and conditions remains challenging due to differences in EVs isolation and quantification methodologies across studies. Nevertheless, the number of EVs recovered during the stationary phase in our experimental setup was quite similar as reported elsewhere, despite apparently lower bacterial growth under these infection-like conditions. N315 EVs exhibited the typical cup-shaped morphology under TEM and carried a complex molecular cargo, including proteins, DNA, RNA, LTA, and PG. As reported for other *S. aureus* EVs, N315 EVs were enriched in specific proteins compared to whole cells, notably lipoproteins, membrane-associated proteins, and cell envelope components (17, 42). We demonstrated that these proteins are integral EV components rather than external contaminants, with a subset localized within the vesicles lumen. Previous studies have shown that EVs produced by human and animal *S. aureus* isolates share a highly conserved core proteome (15, 42, 26). N315 EVs follow this trend, as revealed by LC-MS/MS analysis, and we confirmed the presence of one of the core proteins, enolase (Eno), within the vesicles using a GFP fusion approach. The presence of RNA associated with *S. aureus* EVs has also been previously reported, and here we confirmed their localization inside EVs using confocal microscopy combined with an RNase protection assay (23, 14, 16, 33). RNA profiling revealed a predominance of small RNAs (∼ 100 nucleotides), as described for EVs from various Gram-positive and -negative bacteria, raising questions about their potential biological functions and contribution to EV-mediated host-pathogen interactions (see below) (54, 55, 14, 56). Finally, although fewer studies have focused on these molecules compared to proteins and RNAs, the presence of DNA, LTA and PG associated with *S. aureus* EVs has also been reported (23, 57).

Several components of the molecular cargo of N315 EVs represent potential PAMPs, *i.e.*, bacterial molecules specifically recognized by host pattern recognition receptors (PRRs) as danger signals. Such recognition initiates signaling cascades that lead to the secretion of pro-inflammatory cytokines and the production of defense molecules involved in pathogen elimination (58). PRRs include different classes of receptors, notably TLRs and NLRs, each involved in the recognition of distinct PAMPs. For instance, bacterial lipoproteins are sensed by the TLR2/TLR1 and TLR2/TLR6 heterodimers, RNA by TLR3 and TLR7, and bacterial peptidoglycan fragments by NOD2. Given the diversity of their cargo, N315 EVs may carry ligands capable of activating multiple PRRs, which trigger distinct immune pathways and host responses. Activation of a PRR by its ligands is known to trigger positive or negative feedback loops that regulate receptor expression and downstream signaling (59). We used this autoregulatory property to investigate which PRRs could potentially interact with N315 EVs by using PRR gene expression levels as a proxy for ligand-receptor binding. Our analysis revealed a selective induction of *TLR1*, *TLR2*, *TLR3*, *TLR4*, *TLR6*, *TLR7*, and *NOD2*. Notably, for TLR2, we confirmed that the modulation of its expression by EVs is coupled with an increase in TLR2 protein levels, and associates with the ability of EVs to directly bind the receptor at the cell surface. Supporting our findings, a recent study using similar reporter cell lines showed that EVs secreted by *S. aureus* NCTC 6571 also interact with TLR2, TLR4, TLR7 and NOD2 (23). In addition, the authors reported that EVs also interacted with TLR8 and TLR9, two receptors that recognize RNA and unmethylated CpG DNA motifs, respectively. We were unable to evaluate TLR8 expression in MG-63 cells due to its very low basal transcript levels. Interestingly, although N315 EVs contain DNA, TLR9 expression remained unchanged, suggesting that N315 EVs do not interact with this receptor. Whether unmethylated CpG motifs are absent from N315 EV-associated DNA or inaccessible to TLR9 should be further assessed to investigate the strain-dependent activation of this TLR. Note that we observed an upregulation of STING, a protein involved in the cytosolic detection of double-stranded DNA. STING, which is activated by cGas, a cytosolic sensor of DNA, triggers IRF3 phosphorylation and promotes the production of type I interferons (60). This suggests that EV-associated DNA could bypass the endosomal sensing via TLR9 to be detected through the cytosolic cGAS– STING pathway and then trigger the production of type I interferon. Another intriguing finding is the interaction between EVs from both N315 and NCTC 6571 with TLR4 (23). Although TLR4 is primarily involved in recognizing bacterial lipopolysaccharide (LPS), a specific constituent of cell walls of Gram-negative bacteria, it is also known to sense other ligands such LTA, a component associated with N315 EVs (61). Indeed, Takeuchi *et al*. (1999) demonstrated a highly decrease of IL-6 and NO_2_^−^ responses in macrophages from TLR4-deficient mice after stimulation with *S. aureus* LTA (61). However, the precise nature of components associated to N315 and NCTC 6571 EVs recognized by TLR4 remains to be elucidated.

Our results highlight the ability of EVs to simultaneously activate multiple PRR sensing pathways, in a manner that may be strain-dependent (*e.g*., TLR9 activation), thereby reinforcing the importance of characterizing EVs derived from various strains. PRRs found in the cytoplasm (NOD2), endosomal compartments (TLR3, TLR7 and TLR4), and at the cell surface (TLR1, TLR2, TLR4, and TLR6) are involved in these sensing pathways. This suggests that the corresponding PRR ligands carried by EVs must be accessible to these different cellular compartments. Lipoproteins, which are surface-exposed, can directly interact with extracellular TLRs from the EV surface. In contrast, PG fragments, which are likely exposed at the EV surface, need to be internalized into the cell to engage NOD2. Likewise, RNA contained within EVs requires prior EV internalization into endosomes and subsequent EV lysis in order to activate TLR3 and TLR7. By confocal microscopy and colocalization analysis, we confirmed the presence of intact EVs inside host cells. Furthermore, by using selective inhibitors, we demonstrated that EV uptake occurs primarily via a dynamin-dependent endocytosis pathway, supporting the activation of endosomal PRRs by EVs. When EVs were double-labelled, confocal microscopy revealed a delocalization of EV-derived signals (*i.e*., membrane *vs* RNA) within late endosomes, indicating the release of RNA-associated EV contents into this compartment. This release, likely mediated by EV lysis, is consistent with the activation of endosomal PRRs, such as TLR3 and TLR7, by EV-associated RNA. As mentioned previously, EVs or EV-derived fragments need to reach the cytoplasm to activate NOD2. This raises the question of whether intact EVs can enter cells through alternative uptake mechanisms and what happens to EVs and their cargo beyond the late endosomal stage.

The dynamin-dependent endocytosis has already been identified as potential pathway for the entry of EVs from MRSA inside host cells (27). However, it has been reported that *S. aureus* EVs can also be internalized by cholesterol-dependent fusion with plasma membrane of host cells (28, 29). These findings open the possibility that alternative modes of EV entry may coexist, as has been described for EVs derived from Gram-negative bacteria (62). In line with this hypothesis, while dynasore reduces *TLR3* and *TLR7* expression (likely by limiting EV entry into endosomes and thus TLR ligand availability), *NOD2* remains unaffected, supporting its activation by N315 EVs through an alternative pathway. The downstream fate of EV cargo following its release within late endosomes warrants further investigation, particularly regarding its potential activity within the cytoplasm. Several studies, mainly conducted from EVs secreted by Gram-negative bacteria, have reported that EV-associated RNAs play important roles in host-pathogen interactions, cell-to-cell communication, and bacterial pathogenesis (55, 63, 64). The favored hypothesis is that intravesicular RNAs are delivered into host cells, where they can modulate gene expression in target cells. These RNAs may activate endosomal TLRs (*e.g.,* TLR3 and TLR7) to initiate downstream signaling pathways and immune responses, which is in line with our findinds. Beyond endosomal sensing, growing evidence supports a potential role for EV-associated RNAs in the cytoplasm. For example, *Legionella pneumophila* EVs carry small RNAs that function similarly to miRNAs and, once translocated into host cells, downregulate innate immune response sensor and regulator proteins (64). Similarly, it has been demonstrated that *S. aureus* EV-associated RNAs activate the cytosolic RNA sensors RIG-I and MDA5 to produce stimulatory responses without the involvement of TLR3, TLR7, or TLR8 (65). These results suggest that RNAs carried by N315 EVs can also reach the cytoplasm following their release from endosomes, where they might interact with alternative intracellular sensing pathways. This hypothesis is further supported by previous reports showing that certain *S. aureus* virulence factors (*e.g.*, protein A, alpha-toxin) are efficiently delivered into host cells via EVs (28, 29), suggesting that at least part of the EV cargo can reach the cytoplasmic compartment and exert functional effects.

Upon binding to their ligands, PRRs (*e.g.*, TLR2/TLR1, TLR2/TLR6, TLR3, TLR4, TLR7, NOD2) initiate intracellular signaling cascades through the recruitment of adaptor proteins such as MyD88, TRIF, and RIP2. These adaptor molecules trigger downstream pathways, involving specialized kinases as MAPKs (p38, ERK, JNK) or IKK (IKK-α, IKK-β, IKK-γ). These latter are able to activate transcription factors NF-κB, and interferon regulatory factors (IRFs), leading to the transcription of immune-related genes (58). As a result, host cells produce various effectors, including pro-inflammatory cytokines and chemokines, which coordinate local immune responses and recruit immune cells to sites of infection. Consistent with this, N315 EVs upregulated the expression of key pro-inflammatory cytokines in MG-63 cells such as IL1-β, IL-6, CXCL8, IFN-β and CCL2, in line with previous studies on *S. aureus* EVs (22–26, 66). At the signaling level, increased expression of *NFκB*, and levels of NFκB1 and NFκB2 proteins, confirmed the activation of the NF-κB pathway. Interestingly, the TLR-dependent activation of this pathway also provides the signal required for NRLP3 inflammasome activation and pyroptosis, by inducing expression of *NRLP3* and *IL1β*, a phenomenon previously described in macrophages exposed to *S. aureus* EVs (24, 27). In our study, both were significant induced after 3 hours of EV exposure, while the transcription of *IL18* and *CASP1,* two other key components of the pathway, remained unchanged. At the protein level (Fig. S3), we detected after 24 hours of EV stimulation a slight increase in IL-1 and IL-18 receptor levels, along with an upregulation of ASC, a key adaptor molecule required for inflammasome complex formation (67). *S. aureus* has been described as capable of being internalized into MG-63 cells and inducing inflammasome activation and IL1β secretion (68). This indicates that N315 EVs are able to partially trigger the inflammasome pathway in non-immune cells, regardless of the presence of the bacteria itself. However, additional stimuli or longer exposure may be required for full activation and pyroptosis.

Regarding other signaling pathways, although no detectable activation of *IRF* genes was observed, N315 EVs induced a strong upregulation of *INFB1* expression (over 18-fold), suggesting the activation of IRF-dependent signaling pathways. Moreover, the decrease in *IRF3* and *IRF7* expression when EV internalization was inhibited supports a role of EVs in IRF pathway activation. However, it is established that NF-κB can also regulate *IFNB1* expression despite this is not its primary route (69, 70). Further studies are required to determine whether the induction of *INFB1* expression by EVs is mediated via the NF-κB or IRF signaling pathway.

Additionally, several STAT family proteins, such as STAT2 and STAT1p91, were among the most upregulated proteins. These proteins are key components of the JAK/STAT pathway, which is typically activated by type I interferons (IFN-α/β) and involved in inflammation, immune cell recruitment, survival, and proliferation (71). Interestingly, SOCS6, a negative regulator of this pathway, was slightly downregulated. Since SOCS proteins mediate feedback inhibition to restore homeostasis, both upregulation of STAT proteins and downregulation of SOCS6 suggest that inflammatory signaling remains active 24 hours after EV exposure. While bacterial EVs are well known to modulate host immune responses and promote proinflammatory cytokine release (23, 27–29, 32, 33), their direct involvement in JAK/STAT pathway activation was not described. Our data suggest either an uncharacterized direct activation mechanism or an indirect effect mediated by host-derived cytokines, such as IFN-β, produced in response to N315 EV exposure.

Altogether, our findings highlight the capacity of N315 EVs to interact with diverse PRRs and activate multiple immune signaling pathways in non-immune cells. This supports the ability of EVs to serve as vehicles for the delivery of immunostimulatory ligands across cellular compartments, ultimately modulating host responses during infection. Although most of our observations were obtained *in vitro*, they raise questions regarding the role of EVs during *S. aureus* infections *in vivo*. EVs, by contributing to the early detection of bacteria by the host immune system and amplifying local inflammation, may act as a decoy and facilitate immune evasion of producing bacteria. Further studies are required to determine the fate and function of EVs during the course of infection, their biodistribution within host tissues, and their contribution to virulence or immune modulation. A better understanding of these mechanisms could reveal novel strategies for therapeutic intervention or vaccine development targeting bacterial EVs.

## ACKNOWLEDGMENTS

This work benefited from the facilities and expertise of the MRic-TEM (https://microscopie.univ-rennes.fr/electronic-microscopy-rennes-mric-tem), MRic-Photonics (https://microscopie.univ-rennes.fr/photonic-microscopy-rennes-mric-photonics) and CytomeTRI (https://biosit.univ-rennes.fr/ressources/cytometri#p-815) platforms from Biosit. The authors are grateful to Agnès Burel, Nina Soler and Xavier Pinson for advices and sessions with the microscope, as well as Alexis Aimé and Laurent Deleurme for flow cytometry sessions. The authors thank Aurélie Nicolas (INRAE, STLO) for her helpful advice on microscope image processing, Jordan Ossemond (INRAE, STLO) for his assistance and helpful advice on microscope image acquisition and analysis, and Mathilde Lécot for her technical assistance.

